# Complete vegetative propagation via an epigenetic state change

**DOI:** 10.64898/2026.03.27.714943

**Authors:** Nicole K. Smoot, Yinwei Zeng, Rebecca M. Hochman, Ben P. Williams

**Author notes:** Correspondence to Ben P. Williams.

## Abstract

The plant kingdom exhibits a wide range of phenotypic variation in capacity to regenerate tissues and organs, from whole-plant vegetative propagation via cuttings, to recalcitrance even under optimized tissue culture. Currently, the molecular pathways underpinning this phenotypic variation are poorly understood. Here, we report that Arabidopsis mutants of the DNA demethylase pathway exhibit dramatically enhanced regeneration and the ability to propagate whole plants from cuttings without the use of exogenous hormones. Vegetatively propagated plants possess a shared regeneration signature of de novo DNA methylation gains at the transcription start sites of many genes, including approximately 30 genes involved in cellular pluripotency and tissue regeneration. These methylation changes can be inherited through sexual reproduction and are associated with exacerbated transcriptomic changes. We propose that loss of the DNA demethylase pathway unlocks a path on the epigenetic landscape towards increased pluripotency and regeneration without the application of exogenous hormones.

## Introduction

DNA methylation is an epigenetic modification in eukaryotes that contributes to genome stability, transcriptional regulation, and the regulation of genomic imprinting (Law and Jacobsen 2010). In both plants and animals, cytosine methylation is enriched over transposable elements and other repetitive sequences, where it reinforces transcriptional silencing and protects genome integrity (Williams and Gehring 2020). Methylation can also extend into nearby regulatory regions of coding genes, impacting expression at some, but not all, loci. In plants, DNA methylation occurs in multiple sequence contexts (CG, CHG, and CHH) and is maintained by multiple distinct enzymatic pathways (Law and Jacobsen 2010). Methylation can be established de novo in any sequence context by the RNA-directed DNA methylation (RdDM) pathway (Erdmann and Picard 2020), which can underpin the formation of new epialleles (Weigel and Colot 2012).

The roles of DNA methylation dynamics in the regulation of pluripotency and differentiation appear to be distinct between plant and animal lineages. In mammals, dynamic changes to DNA methylation accompany major developmental transitions. During early embryogenesis and the exit from pluripotency, de novo methyltransferases and maintenance methyltransferases establish lineage-specific methylation patterns, and Ten–eleven translocation (TET) family demethylases actively erase methylation at key regulatory loci (Bagci and Fisher 2013; Seisenberger et al. 2013). Studies in mice show that TET activity is critical for development and its disruption prevents a proper exit from pluripotency and causes failure to establish differentiated cell fates. As a result, embryos display severe developmental defects and some perinatal lethality (Dawlaty et al. 2013; Kang et al. 2015), (Dawlaty et al. 2014). In plants, however, the importance of DNA methylation dynamics in regulating somatic cell identity and pluripotency is less clear. Active removal of methylation in plants is performed by an evolutionarily distinct mechanism, instead mediated by a family of 5-methylcytosine DNA glycosylases, DEMETER (DME) (Choi et al. 2002), REPRESSOR OF SILENCING 1 (ROS1) (Gong et al. 2002), and DEMETER-LIKE 2 and 3 (Penterman et al. 2007), collectively named DRDD. Genetic studies have shown that DNA demethylation is indispensable for reproductive development: DRDD protein family activity in the central cell and endosperm is required for genomic imprinting and proper seed development in Arabidopsis (Choi et al. 2002), maize (Gent et al. 2022), and rice (Ono et al. 2012), and demethylation in male gametophytes is required for normal pollen function (Khouider et al. 2021; Zeng et al. 2024). By contrast, the somatic tissues of DNA demethylase mutants are relatively normal at the macroscopic level, with most tissues and organs appearing to possess a typical organization and arrangement of cellular identities (Williams et al. 2022). However, closer examination at cellular resolution has identified subtle developmental defects. In the shoot epidermis, demethylase mutants exhibit overproliferation of stomatal precursor cells (Yamamuro et al. 2014; Kim et al. 2021) and in vascular tissues, demethylase mutants show a failure to develop unbroken tracheary elements (Lin et al. 2020). Additionally, roots of homozygous *dme* mutants show altered organization of the meristem (Kim et al. 2021). These phenotypes demonstrate that active DNA demethylation in plants does not broadly influence pluripotency and the core identity of most cell types as it does in mammals, but nevertheless can influence developmental traits if genes underpinning those traits are demethylase targets (Yamamuro et al. 2014; Williams et al. 2022).

The plant kingdom possesses an extraordinary capacity for cellular reprogramming and regeneration. Differentiated cells can acquire pluripotency and regenerate entire organs or even whole plants under appropriate conditions. This developmental plasticity underpins genetic transformation, tissue culture and clonal propagation. Both DNA methylation and histone modification pathways have been implicated as important in the regulation of plant regeneration (Shemer et al. 2015; Liu et al. 2018; Ishihara et al. 2019; Rymen et al. 2019; Temman et al. 2023), yet the precise roles of epigenetic pathways in regeneration processes are poorly understood. Loss-of-function mutants of the DNA demethylase DME exhibit enhanced regeneration under tissue culture, suggesting a role for this pathway in modulating pluripotency and regeneration (Kim et al. 2021). Given the central role of DNA methylation dynamics in controlling the exit from pluripotency in animals, whether active DNA demethylation modulates pluripotency networks and regenerative competence in plant somatic tissues is a pertinent question.

Here, we address this question by interrogating the consequences of disrupting the DNA demethylase pathway on cell identity and regenerative potential in Arabidopsis. We show that mutants of the DNA demethylase pathway in Arabidopsis display dramatically enhanced regeneration abilities, even without the application of exogenous hormones, effectively enabling vegetative propagation from cuttings. Regenerated demethylase mutant plants also possess a unique set of regeneration-associated DNA methylation changes, directly impacting a number of pluripotency and regeneration genes.

## Results

### DNA demethylation represses organ regeneration with and without callus induction

To define the functional significance of DNA demethylation in somatic development in Arabidopsis, we previously generated quadruple somatic *drdd* mutants (Williams et al. 2022), which carry homozygous loss of function alleles for each of the four DNA demethylase genes. Our previous RNA-seq analysis of *drdd* rosette leaves identified a pattern of upregulation of a number of root meristem and shoot apical meristem-associated genes that are normally silent or lowly expressed in shoot tissues (Williams et al. 2022), which has also been independently described in homozygous *dme* mutants (Kim et al. 2021; Lee et al. 2024). To investigate these transcriptome changes in more depth, we expanded our RNA-seq profiling of *drdd* shoot tissues to include seven biological replicates per genotype from wild-type (WT, a closely related segregate from *drdd* heterozygotes (Williams et al. 2022)) and *drdd* mutants. We identified 762 differentially expressed genes (DEGs) (Dataset S1), increasing our previous estimate of transcriptome differences between *drdd* and WT by more than 50%. Among these DEGs was *WUSCHEL-HOMEOBOX 5* (*WOX5*), the root meristem and pluripotency factor that is normally not expressed in mature leaf tissues. We also detected elevated expression of three other WOX family genes important for cellular pluripotency and regeneration (Liu et al. 2014; Lee et al. 2022)(Fig. 1A). We validated the increased expression of *WOX5* and *WOX11* using qRT-PCR and found both to be strongly upregulated (5-60-fold) in mature leaf tissues of *drdd* mutants (Fig. 1B).

**Figure 1.**
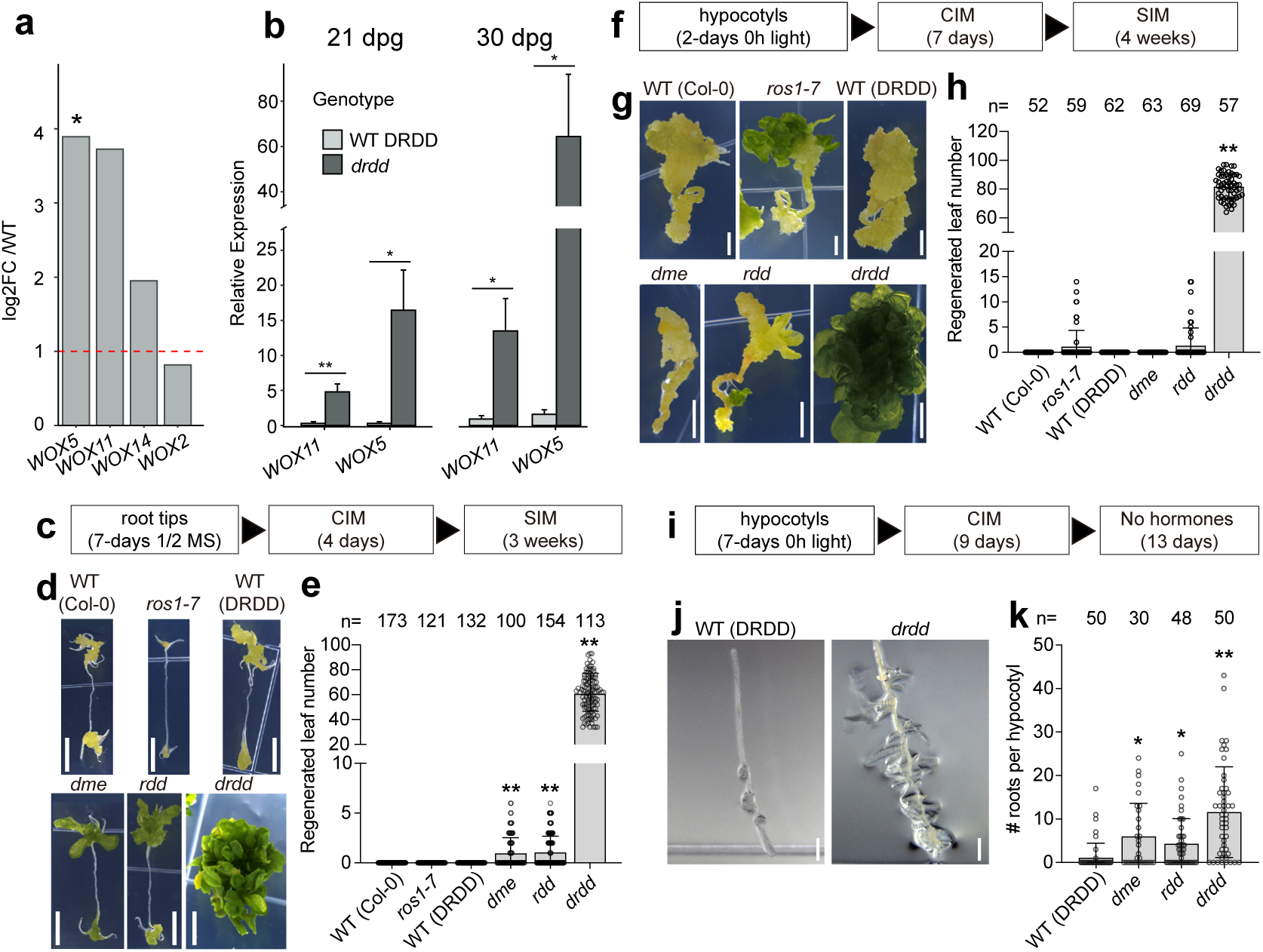
DNA demethylase mutants show increased regenerative capacity across multiple tissues. A) RNA-seq data showing fold change of *WOX* genes in *drdd* leaf tissue relative to WT expression levels. *=padj<0.05. B) Relative expression in WT and *drdd* from qRT-PCR. Tissue is first true leaves from 21 and 30dpg plants. Expression is normalized to WT expression of each gene. Significance calculated with T test. *=p<0.05, ** = p<0.01. C) Summary of tissue culture regimen for root tip > shoot regeneration assay. CIM = callus induction medium, SIM = shoot induction medium. D) Representative images of cultured root tips after tissue culture as specified in C. Scale bar = 5 mm. E) Number of regenerated leaves per root tip explant, n = number of individual explants assayed for each genotype. Statistical analysis: Kruskal–Wallis test (H(5)=569.9, p<0.0001) with Dunn’s multiple-comparisons test (mutant vs appropriate WT: WT[Col-0] vs *ros1*, WT[DRDD] vs *dme*, *rdd* and *drdd*), **p<0.0001. F) Summary of tissue culture regimen for hypocotyl > shoot regeneration assay. G) Representative images of cultured hypocotyl explants after tissue culture as specified in F. Scale bars = 2 mm. H) Number of regenerated leaves per hypocotyl explant, n = number of individual explants assayed for each genotype. Statistical analysis: Kruskal–Wallis test (H(5)=298.3, p<0.0001) with Dunn’s multiple-comparisons test (mutant vs appropriate WT), **p<0.0001. I) Summary of tissue culture regimen for hypocotyl > adventitious root regeneration assay. J) Representative images of hypocotyls after tissue culture as specified in I. K) Number of adventitious roots per hypocotyl explant, n = number of individual explants per genotype. Statistical analysis: Kruskal–Wallis test (H(3)=52.56, p<0.0001) with Dunn’s multiple-comparisons test (mutant vs WT), *p<0.01, **p<0.0001. In B, E, H and K, Bars represent mean values and error bars represent standard deviation.

As both WOX5 and WOX11 function in organ regeneration (Sugimoto et al. 2010; Liu et al. 2014; Kim et al. 2018; Lee et al. 2022), and DNA methylation and demethylation pathways have previously been implicated in the regulation of plant regeneration (Shemer et al. 2015; Liu et al. 2018; Kim et al. 2021), we therefore sought to examine whether *drdd* mutants exhibit enhanced regeneration capacity. We first tested shoot regeneration from root tip explants under a callus induction medium (CIM) > shoot induction medium (SIM) tissue culture protocol (Fig. 1C). In this assay, *drdd* mutants exhibited dramatically enhanced shoot regeneration compared to WT, which did not regenerate under these conditions (Fig. 1D-E, Table S2). Both *dme* (Williams et al. 2022) and *rdd* (Penterman et al. 2007) (weaker mutants that do not share genetic ancestry) exhibited a weaker version of this phenotype (Fig. 1D-E, Table S2), strongly suggesting that enhanced regeneration in *drdd* is due to loss of DNA demethylase activity and not caused by secondary mutations linked to *drdd*. Additionally, we found that etiolated hypocotyl explants of *drdd* mutants also exhibited enhanced shoot regeneration compared to WT (Fig. 1F-H, Table S2). *ros1-7* (Williams et al. 2015) and *rdd* mutants (which do not share genetic ancestry) also exhibited enhanced regeneration from hypocotyls compared to WT, albeit to a lesser extent.

To investigate whether DNA demethylation mutations also affect root organogenesis, we performed an assay in which etiolated hypocotyls were first incubated on CIM then transferred to hormone-free medium to induce adventitious root regeneration (Fig. 1I). *drdd* mutants exhibited substantially higher rates of root organogenesis compared to WT, with *dme* and *rdd* also exhibiting increased root regeneration (Fig. 1J-K, Table S2). We even observed a high frequency of adventitious root organogenesis in *drdd* hypocotyls incubated under hormone-free conditions, with 63% of hypocotyls regenerating at least one adventitious root, compared to only 3% of WT (Figure S1, Table S2). Together these data suggest that loss of DNA demethylase function in somatic cells leads to substantially enhanced capacity of both shoot and root organogenesis, even in conditions where regeneration from WT explants is minimal.

To further examine the shoot regenerative capacity of DNA demethylation mutants, we also performed a less hormonally intensive regeneration assay, in which hypocotyls were incubated on hormone-free growth medium (Fig. 2A), followed by transfer to SIM. Under these conditions, WT Arabidopsis plants are unable to regenerate (0% shoot regeneration, n = 143, Table S2). However, a subset of *drdd* mutant hypocotyls exhibited shoot regeneration (10% shoot regeneration, n = 145), despite the fact that conventional callus induction was not performed (Fig. 2B-C). Notably, we also observed similarly enhanced shoot regeneration in *dme* single mutants and triple *rdd* mutants (Fig. 2B-C, Table S2), providing confirmation of this phenotype across multiple independent mutant backgrounds.

**Figure 2.**
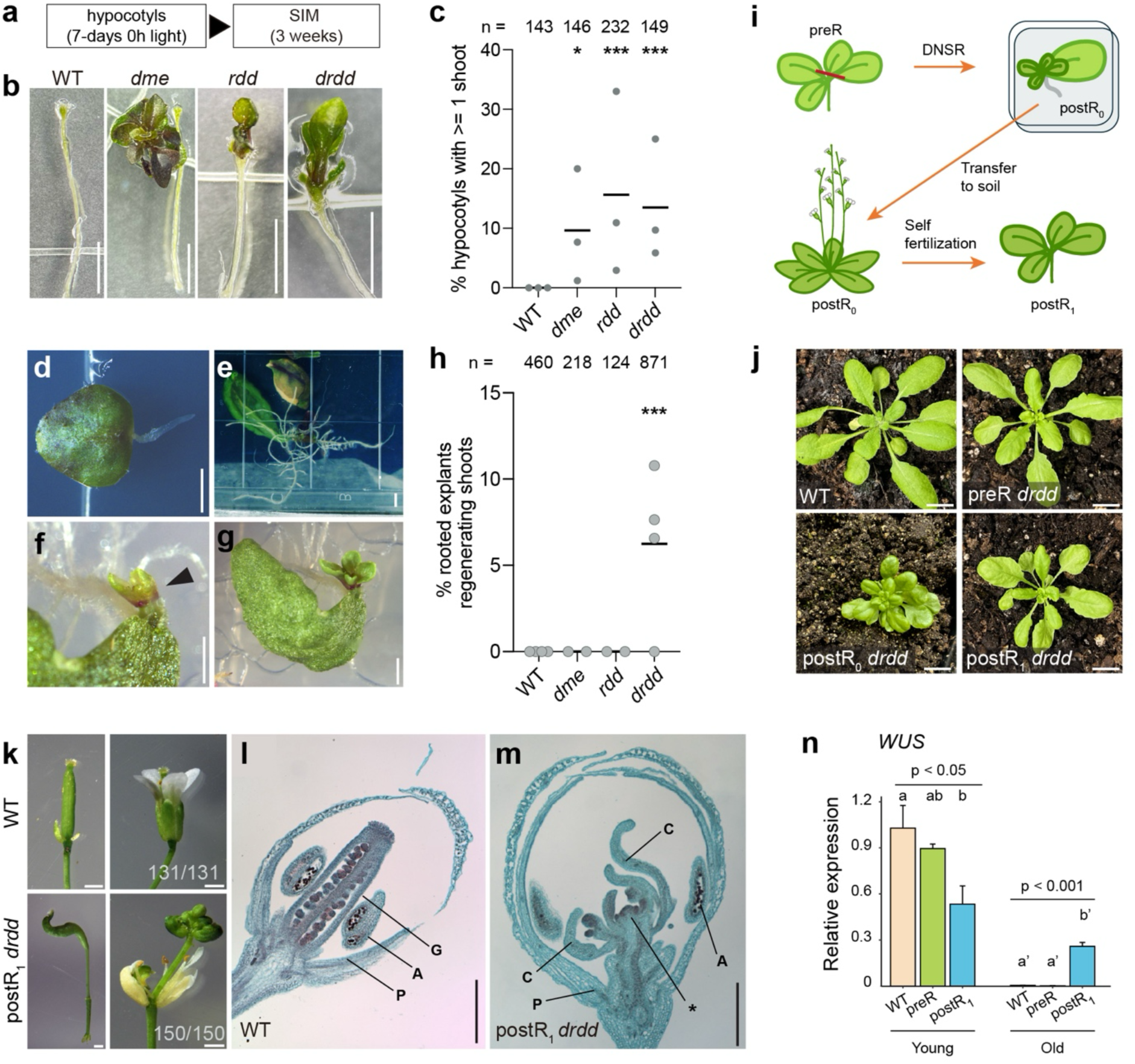
*drdd* mutants are capable of whole plant propagation without exogenous hormone application. (A) CIM-free hypocotyl-to-shoot regeneration protocol. (B) Representative images of hypocotyl explants after the protocol in (A). (C) Percentage of explants regenerating shoots. N, independent experiments; n, total explants across experiments. (D, E) Representative images of *drdd* leaf explants after 8 d (D) and 2 weeks (E) on hormone-free Gamborg B5 medium. (F, G) Representative images of shoot-regenerating *drdd* leaf explants after 4–6 weeks; the arrow indicates a shoot meristem. (H) Percentage of *drdd* leaf explants that spontaneously regenerated shoots on hormone-free medium; n indicates total explants across experiments. In (C) and (H), horizontal lines indicate medians, boxes indicate the 25th–75th percentiles, and whiskers indicate minima and maxima. Significance was assessed using Fisher’s exact test: *P < 0.01; ***P < 0.0001. (I) Whole-plant vegetative propagation workflow. DNRR, de novo root regeneration; DNSR, de novo shoot regeneration; preR, non-regenerated *drdd*; postR_0_, regenerated *drdd*; postR_1_, selfed, seed-derived progeny of postR_0_. (J) Representative vegetatively propagated *drdd* plant generated using the workflow in (I). (K) Representative images of dissected gynoecia (left) and flowers (right) from WT and postR_1_ *drdd* plants. Numbers indicate plants exhibiting the phenotype among all plants examined. (L, M) Transverse sections of mature WT (L) and postR_1_ *drdd* (M) flower buds. G, gynoecium; C, carpel; A, anther; P, petal; asterisk, unterminated meristem. (N) WUS expression measured by qRT-PCR in young (stages 6–9) and old (stages 9–12) floral buds and normalized to young WT buds. Significance was assessed by ANOVA followed by Tukey’s HSD test. Scale bars: 2 mm in (B) and (D–G); 1 mm in (K); 250 μm in (L, M).

### Enhanced regeneration of *drdd* mutants enables complete vegetative propagation without exogenous hormones

We also tested whether DNA demethylation mutants exhibit enhanced de novo root organogenesis from leaf cuttings in the absence of exogenous hormones, as previously described (Liu et al. 2014). Under these conditions, first true leaves were excised and placed on hormone-free medium, we observed comparable root organogenesis rates between WT and *drdd* mutants (Figure S1). However, while performing this experiment, we noticed that *drdd* leaf cuttings exhibited enhanced vigor and survival on hormone-free medium compared to WT (Fig. 2D-E, Figure S2), and a subset of *drdd* explants spontaneously regenerated shoot meristems at the original wound site (Fig. 2F-H, Figure S2). Explants with regenerated shoots were transferred to soil and grown under conventional growth conditions to complete the life cycle (Fig. 2I-J). The spontaneous production of shoot meristems therefore permitted full vegetative propagation of *drdd* mutants, akin to plant species that can be propagated from cuttings. To our knowledge, this is the first example of vegetative propagation from cuttings in Arabidopsis without the use of exogenous hormones or transgenes designed to enhance somatic regeneration or embryogenesis.

Regenerated *drdd* plants (hereafter termed postR_0_) flowered and set seeds normally, permitting further study of the progeny of plants (which we term the postR_1_ generation) that have been through the regeneration process (Fig. 2I). 100% of postR_0_ plants exhibited a striking floral defect in which all of the flowers observed possessed non-terminating meristems after carpel differentiation (Fig. 2K-M). Instead of forming a single gynoecium at the center of the flower, the gynoecium is displaced by aberrant inflorescence meristems, leading to ectopic shoots emerging from flowers (Fig. 2K,M). The phenotype was stably inherited in the progeny of these plants: all observed postR_1_ plants also showed improper meristem termination in all flowers. This mutant phenotype, termed “floral reversion”, has been shown in homeotic floral mutants such as *ap1* (Bowman et al. 1993)*, ag* (Okamuro et al. 1996), as well as the *knuckles* mutant (Payne et al. 2004). We did not observe changes to the identity or number of floral organs, nor did we observe this phenotype in any non-regenerated (preR) *drdd* plants. Consistent with the non-terminating meristem phenotype, we detected transcripts of the meristem identity gene WUSCHEL in stage 9-12 flower buds (Smyth et al. 1990) of postR_1_ *drdd*, but not WT or preR *drdd* (Figure 2N).

### Regenerative propagation of *drdd* is associated with heritable de novo DNA methylation gains

As only a subset of *drdd* explants regenerated whole plants, we hypothesized that these individual regenerated plants could share DNA methylation changes linked to their regenerated state (i.e. not present in non-regenerated *drdd* plants), which may not be detectable in preR *drdd* methylomes. To test this, we performed enzymatic methyl sequencing (EM-seq) on four independent regenerated postR_0_ plants, which we termed postR lines (L1–L4). We then identified differentially methylated regions (DMRs) between non-regenerated and regenerated plants (see methods) (Fig. 3A, Datasets S2, S3 and S4). Each individual postR_0_ plant sequenced possessed a set of approximately 2,000 new CG DMRs that were absent in preR *drdd* mutants (Fig. 3B), which we term regeneration-associated DMRs (R-DMRs). Like preR *drdd* DMRs, regenerated *drdd* DMRs are predominantly located within gene-rich chromosome arms (Figure S3), where active demethylation by DRDD is known to occur (Penterman et al. 2007; Williams et al. 2022). We then sought to determine if independent regenerated plants gain R-DMRs at the same loci, reflecting a possible “regeneration signature” in the methylome, as opposed to random methylation gains in independent postR plants. R-DMRs were highly consistent between lines: 77-85% of DMRs called in any individual line were also called in at least one other regenerated plant (Figure S4). Additionally, the DMRs not called in multiple lines still exhibited increased methylation in all lines, despite being outside the statistical cutoffs to be called as DMRs (Figure S4). Independently regenerated plants therefore exhibit similar methylation profiles.

**Fig 3:**
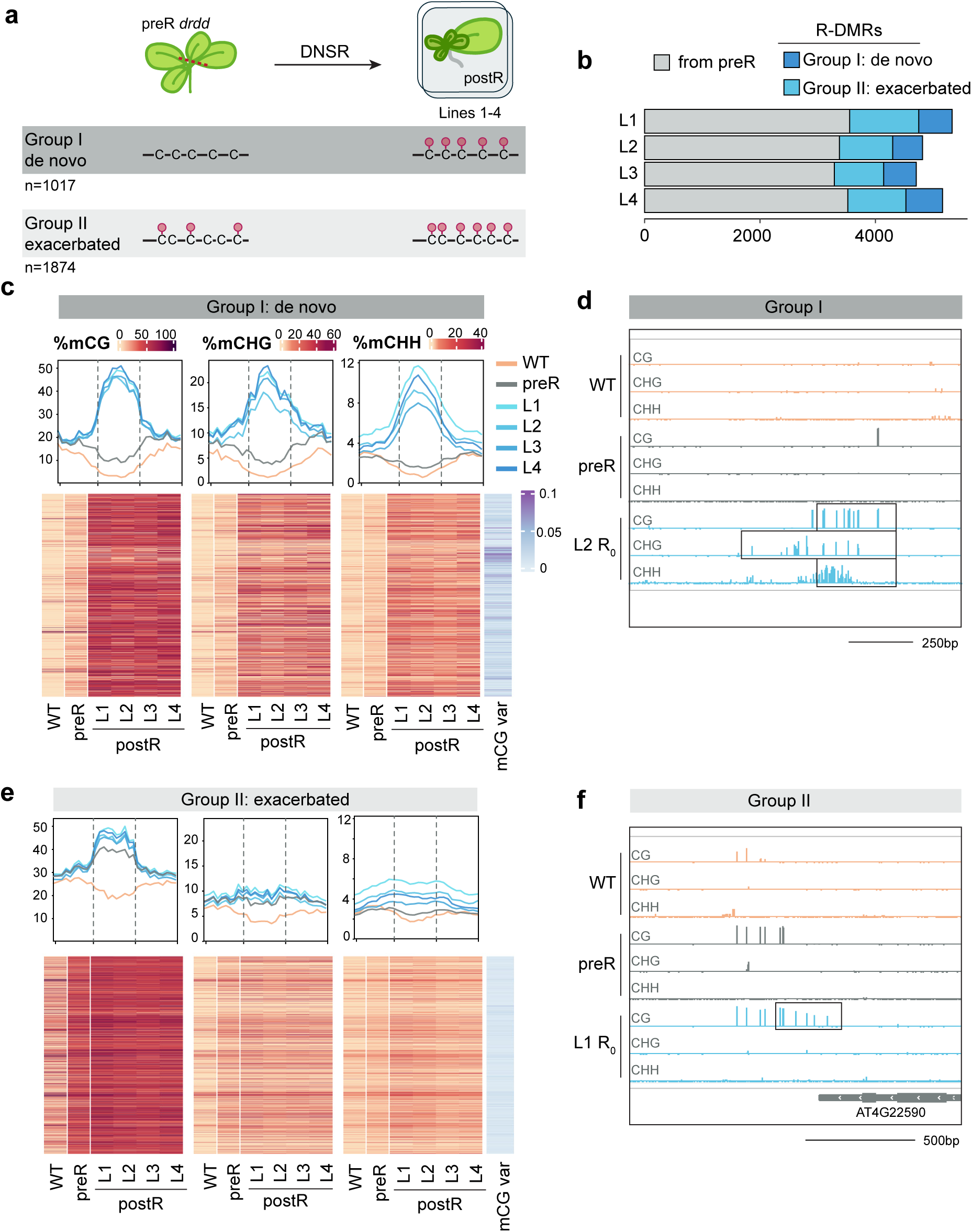
postR drdd methylomes show multicontext regeneration-associated methylation gains. A) DMRs relative to WT gained during regeneration (R-DMRs) in postR *drdd* methylomes were separated into two categories. B) Numbers of DMRs relative to WT in each independently regenerated line (L1-4), broken into categories of inherited with no changes from preR; Group I; and Group II. C) Metaplots and heatmaps of averaged cytosine methylation across de novo DMRs. Dashed lines indicate scaled DMR regions; outer bounds represent 300 bp upstream and downstream of the DMR region. CG methylation variance between regenerated lines is shown on the right. D) Representative screenshot of Group I de novo methylation gain (1:23,365,081-23,366,227). Black box indicates called DMR region. All tracks set to (-10, 100). E) Metaplots and heatmaps of averaged cytosine methylation across exacerbated DMRs. Dashed lines indicate scaled DMR regions; outer bounds represent 300 bp up and downstream of the DMR region. CG methylation variance between regenerated lines is shown on the right. F) Representative screenshot of Group II exacerbated methylation gain. (4:11,892,841-11,894,242). Black box indicates called DMR region. All tracks set to (-10, 100).

To further understand the nature of DNA methylation changes in postR *drdd* plants, we classified them into two groups based on their methylation state in preR *drdd* (Fig. 3A-B, see methods). The first group (Group I -- 1,017 DMRs across all lines) represents regions that are unmethylated in *drdd* and gain de novo methylation after regeneration. These DMRs displayed strong gains in CG, CHG and CHH contexts, indicative of de novo methylation by the RdDM pathway (Fig. 3C-D). Additionally, these DMRs were similarly highly methylated in all four postR plants sequenced, strongly suggesting they represent a regeneration signature of de novo methylation at shared loci. The second group (Group II -- 1,874 DMRs across all lines) was comprised of regions already partially methylated in preR *drdd* (Fig. 3E-F). These regions exhibit partial or slightly exacerbated methylation gains in all three contexts after regeneration. Unlike group I DMRs, group II DMRs exhibit moderate levels of CHG and CHH methylation in preR *drdd* and are therefore likely to be RdDM targets that were already present before regeneration. During regeneration these regions appear to accumulate increased methylation, perhaps due to feed-forward loops that reinforce RdDM in the absence of DRDD activity.

Next, we sought to determine if R-DMRs are meiotically heritable, or if the methylation signature associated with regeneration is specific to regenerated (postR_0_) plants. To test this, we sequenced the methylomes of self-fertilized progeny (postR_1_) from two postR_0_ plants, line 2 and line 4 (L2 and L4 in Figure 3). First, we examined the level of methylation at group I (de novo) and group II (exacerbated) DMRs. At group I DMRs, all three sequence contexts maintained consistently high methylation in the postR_1_ generation (Fig. 4A), indicating that these methylation gains are heritable across generations. In contrast, group II DMRs maintained elevated CG methylation in postR_1_ progeny, but CHG and CHH levels reverted to those already present in preR *drdd* plants (Fig. 4B). To further verify this inheritance pattern, we also called DMRs between postR_1_ *drdd* plants and WT and overlapped these DMRs with group I and group II DMRs from the previous generation. Any DMRs independently identified in both the parent and progeny generations were classified as inherited. With this approach, we observed that the vast majority of group I CG DMRs were inherited (92 and 95% across the two lines), as well as a high percentage of CHG DMRs (57 and 66%) (Fig. 4C). Inheritance was less consistent among CHH DMRs, showing two inheritance patterns: 1) CMT2-type (Gouil and Baulcombe 2016) CHH DMRs in pericentromeric regions were absent in the R_1_ generation (suggesting they are specific to postR_0_ plants, perhaps due to different growth conditions used in the regeneration assay) and 2) RdDM-type CHH DMRs in the chromosome arms were mostly inherited (Fig. 4C, E, Figure S5). In contrast, group II DMRs in each context were less likely to be inherited than their group I counterparts. Overall, these data suggest that once initiated during the regeneration process, new sites of de novo methylation are permanent in both regenerated plants and their progeny.

**Fig 4:**
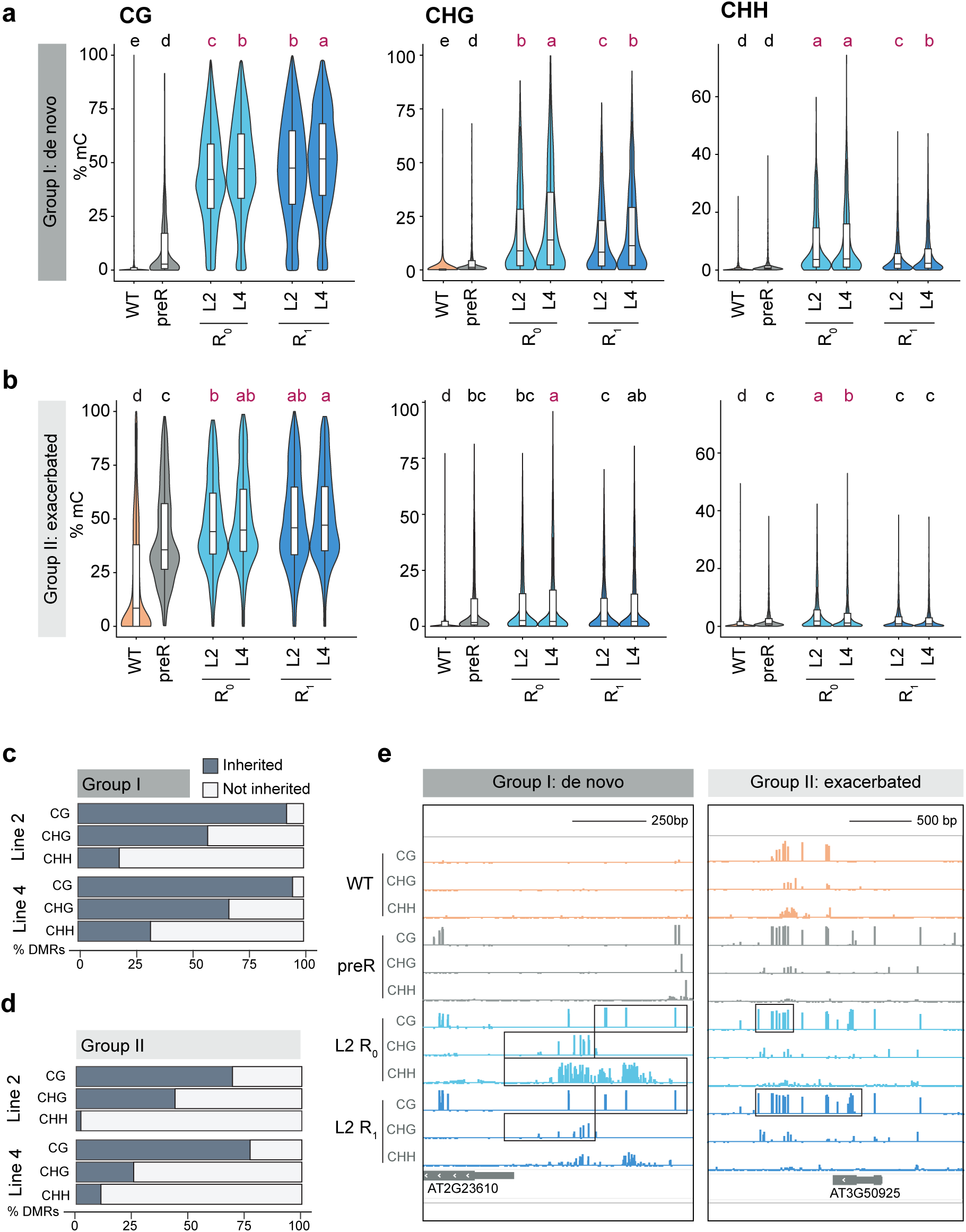
A regeneration signature in the methylome is inherited to progeny. A) Average methylation level of Group I DMRs across all lines in WT, preR *drdd*, postR_0_ lines 2 and 4, and postR_1_ lines 2 and 4. Significance calculated by ANOVA and Tukey’s HSD. Red letters indicate significant difference from preR methylation levels. B) Average methylation level of Group II DMRs across all regenerated lines in WT, preR *drdd*, postR_0_ lines 2 and 4, and postR_1_ lines 2 and 4. Significance calculated by ANOVA and Tukey’s HSD. C) Percentage of Group I de novo DMRs of each context that were also called in the R_1_ generation of each line. D) Percent of Group II exacerbated DMRs of each context that were also called in the R_1_ generation of each line. E) Representative screenshot of Group I de novo DMR inheritance (2:10,046,038-10,047,124). Boxes indicate called DMR region. All tracks set to (-10, 100). F) Representative screenshot of Group II exacerbated DMR inheritance. (3:18,924,698-18,926,466). Box indicates called DMR region. All tracks set to (-10, 100).

### Regenerated *drdd* mutants exhibit exacerbated transcriptomes

We noticed that de novo methylation gains in regenerated plants were frequently in close proximity to the transcription start site (TSS) of genes (e.g., Figure 4E). To quantify this, we plotted the distance of all group I DMRs to the TSS of their nearest gene (Fig. 5A). This revealed a very strong and statistically significant association between group I DMRs and TSSs compared to both randomly selected windows and group II DMRs, which showed a weaker association (Fig. 5A). To examine the impact of these TSS-associated R-DMRs on gene expression, we profiled the transcriptome of postR_1_ progeny plants, preR *drdd* and WT controls, using seven independent biological replicates per genotype. We reasoned that postR_1_ plants were the best comparison to non-regenerated plants, as they could be grown in identical growth conditions with multiple biological replicates, as opposed to immediately regenerated individual plants that were incubated on growth medium for several weeks. Furthermore, the majority of group I DMRs are faithfully inherited in postR_1_ plants (Fig. 4C-D, Figure S5). When compared to WT, postR_1_ *drdd* exhibited 1,363 differentially expressed genes (DEGs), almost double the number between preR *drdd* vs WT (762 genes) (Fig. 5B, Dataset S5). The majority of DEGs in preR *drdd* were also differentially expressed in postR_1_ *drdd* (Fig. 5C), but regenerated-plant progeny gained hundreds of new DEGs as a consequence of increased transcriptome divergence from WT plants (Fig. 5B-C, Figure S6). Consistent with this, postR_1_ *drdd* transcriptomes showed a lower correlation with WT than did preR *drdd* transcriptomes (Figure S6). Regeneration therefore exacerbates the transcriptional differences of demethylase mutants and is associated with the gain of hundreds of DEGs.

**Fig 5:**
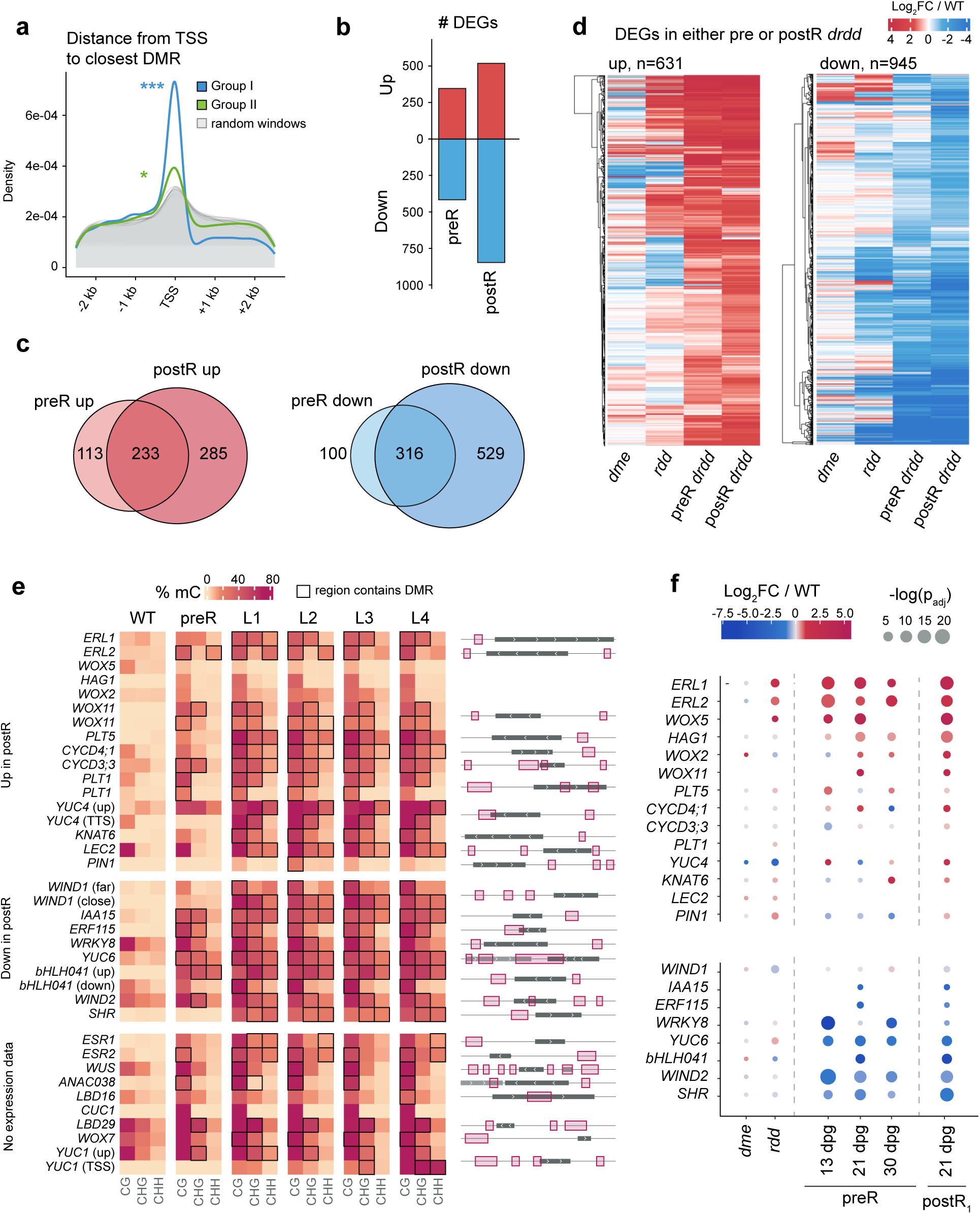
postR drdd shows an exacerbated transcriptional divergence from preR. A) Distance from transcription start sites to Group I and II DMRs. Controls are a set of random windows equivalent to the number of DMRs in each category. Significance calculated by K-S test. Group I vs random windows: p=0.000149 and D=0.0169. Group II: p=0.01215 and D=0.0124. B) Number of differentially expressed genes (DEGs) relative to WT in preR *drdd* and postR. C) Overlap of DEGs between preR and postR. D) Expression level relative to a corresponding WT of all preR and postR DEGs across *dme*, *rdd*, preR *drdd*, and postR *drdd*. Each genotype was normalized to its own WT control sample. E) Heatmap showing average methylation level in each sequence context at regions adjacent to or overlapping with genes involved in plant tissue regeneration (region coordinates found in Extended Data Table 8). Black outlines denote samples in which a statistically significant DMR was called at each locus. Genes are grouped by the direction of their expression change in postR_1_ *drdd* vs WT. Selected regions for heatmap are within 2.5 kb of the gene, and non-overlapping with any other genes. Diagrams of genes with DMRs indicate all proximal differentially methylated regions (pink boxes) relative to gene body (dark grey boxes), and any neighboring genes (light grey boxes). F) Expression changes of regeneration-associated genes in *dme*, *rdd*, preR *drdd* and postR_1_ *drdd*. Fold change is relative to a corresponding WT sample grown in the same conditions. No expression was detected for *ESR1*, *ESR2*, *WUS*, *ANAC038*, *LBD16*, *CUC1*, *LBD29*, *YUC1*, or *WOX7*.

We reasoned that R-DMR gains at or near TSSs could contribute to transcriptomic differences between preR and postR_1_ *drdd,* including expression changes that could underpin the enhanced regeneration phenotypes of regenerated *drdd* lines. We therefore examined the methylation profiles of 31 genes known to be involved in one or more regeneration pathways based on previous literature (Fig. 5E; Table S3, with references therein). Twenty-seven of these genes were within 2 kb of a DMR present in at least one postR0 line, with no intervening genes (Fig. 5E), indicating that the regeneration signature of methylation changes broadly affects regeneration-associated genes. We examined the expression profiles of these regeneration-associated genes in preR and postR_1_ *drdd* plants of multiple ages, as well as in *dme* and *rdd* (Williams et al. 2022) mutants, which exhibit subtler regeneration phenotypes and more moderate transcriptomic differences (Fig. 5D–F). Of the 22 regeneration-associated genes detected in leaf tissue, 13 differed by at least twofold in postR_1_ *drdd* relative to WT, eight of which were significant (adjusted P value < 0.01, Fig. 5F). For example, both the regeneration activator *WOX11* (Liu et al. 2014) and the recently described regeneration suppressor *bHLH041* (Xu et al. 2024) were significantly up- and downregulated, respectively, and both possessed multicontext DMRs in both their 5’ and 3’ intergenic regions (Fig. 5E-F). Notably, the partial differential expression of regeneration-associated genes in lower order mutants *dme* and *rdd* (Fig. 5F) was consistent with the subtler regeneration phenotypes of these mutants (Figs. 1 and 2). We therefore propose that the extent of DRDD disruption determines the scale of methylome divergence at regeneration-associated genes, and consequently alters the expression profiles of this pathway in somatic tissues. Vegetative regeneration of *drdd* causes these mutants to move further along this epigenetic landscape, perhaps through the accumulation of new DMRs that build up over successive somatic cell divisions or are selected through a cellular bottleneck during the regeneration process.

## Discussion

In this work, we establish that mutants of the DNA demethylase pathway in Arabidopsis exhibit altered expression of pluripotency and regeneration factors, which correspond to enhanced regenerative capacity under conventional tissue-culture conditions (which has also been shown for single *dme* mutants (Kim et al. 2021; Lee et al. 2024)), and in plant-regeneration assays without exogenously supplied hormones. The most extreme expression changes and phenotypes are in the quadruple somatic *drdd* mutants, which are capable of whole-plant vegetative propagation – a trait that is widespread across the plant kingdom but not (to our knowledge) previously reported in *Arabidopsis*. The upregulation of *WOX* pluripotency factors that are also associated with callus induction in *drdd* shoot tissues (Fig. 1A) may explain why callus induction using CIM is dispensable in these mutants.

Vegetatively propagated *drdd* plants exhibited widespread non-random gains in methylation and greater transcriptomic divergence than non-regenerated *drdd*. We reason that two possible mechanisms could underlie this finding. First, DNA methylation gains could accumulate throughout somatic growth and development, leading to exacerbated methylomes in some somatic cells and tissues. Second, the regeneration of *drdd* from cuttings likely occurs through a cellular bottleneck, and this bottleneck could select for “exacerbated *drdd*” epigenetic landscapes in the cells that successfully become founder cells of regenerated plants. We view this hypothesis as plausible, as all regenerated *drdd* plants and their progeny exhibit substantial methylation changes at ∼30 genes known to be important for regeneration in plants (Figure 5E). Additionally, two studies have shown that *Arabidopsis* plants that are clonally propagated via the overexpression of embryogenic transcription programs directly inherit methylation imprints of the founder cells and tissues from which they are regenerated (Wibowo et al. 2018, 2022), further suggesting that the methylomes of regenerated plants could represent the epigenetic profiles of founder cells. We also observed that lower order demethylase mutants *dme* and *rdd* displayed more moderate transcriptome differences and regeneration phenotypes akin to a dosage effect. The extent of DNA demethylase disruption therefore dictates the extent of transcriptome divergence, either through mutation dosage, or a cellular bottleneck in somatic near-knockouts.

In mammals, the TET demethylases are critical for establishing exit from pluripotency and the identity of differentiated cell types (Dawlaty et al. 2013; Kang et al. 2015). In plants, the relatively normal organization of cell types and tissues in most DNA methyltransferase and demethylase mutants has led to a consensus that DNA methylation patterns are not critical for establishing cell identities in plants. However, our study joins a number of previous studies that implicate DNA methylation patterns and dynamics in impacting the flexibility of cell identities and pluripotency during tissue regeneration. Enhanced regeneration has now been demonstrated in relatively severe DNA methyltransferase mutants (Shemer et al. 2015; Liu et al. 2018) and demethylase mutants (Kim et al. 2021; Lee et al. 2024), this study). Additionally, multiple histone modification pathways that interact with the DNA methylation landscape are also important in the regulation of cell identity and regeneration pathways (Ikeuchi et al. 2015; Ishihara et al. 2019; Temman et al. 2023; Rajabhoj et al. 2024). In particular, the removal of H3K4me2 by LDL3 is important for shoot regeneration competence (Ishihara et al. 2019). H3K4me2 overlaps a number of DRDD targets (Williams et al. 2023) and the methylation of H3K4 is required for DRDD recruitment to target loci (Wang et al. 2025). It is possible that H3K4me2 removal promotes regeneration by excluding DRDD activity, which our mutant phenotypes suggest broadly suppresses regeneration pathways.

Regenerated *drdd* plants possess a large number of R-DMRs that plausibly cause transcriptional changes that underpin the dramatic regeneration phenotypes we observe. Currently, the pleiotropy of the *drdd* mutant and the large number of R-DMRs associated with regeneration-regulating genes (Fig. 5E) make further dissection of the mechanism underpinning enhanced regeneration challenging. It is unclear whether the altered regulation of an upstream regulator or the cumulative effect of dozens of DNA methylation changes are sufficient to explain the enhanced regeneration of *drdd*. An additional limitation of our study is that it does not take into account the complex regulatory dynamics that occur within the first few hours and days of the regeneration process (Iwase et al. 2017; Liu et al. 2022), including the wound response, hormone signaling, and the dedifferentiation and proliferation of founder cells. We anticipate that more detailed study of each of these processes in isolation will lead to a more precise mechanistic understanding of the interplay between the DNA methylation landscape and broader regeneration processes. Our study also found that regenerated *drdd* plants exhibited an unusual floral phenotype in which meristems do not terminate upon flower formation (Fig. 2M). It is possible that this phenotype is linked to the enhanced regeneration phenotype of regenerated *drdd* plants and is part of a general programmatic shift towards enhanced pluripotency. While we currently lack evidence to support this claim, we note that other studies of demethylase mutants have identified additional phenotypes consistent with a weakening of fixed cell identities and a tendency towards increased proliferation of more pluripotent cell types (Yamamuro et al. 2014; Iwase et al. 2017; Kim et al. 2021; Liu et al. 2022). Understanding if there is a link between DNA demethylation and a global tuning of pluripotency in all tissues would be an interesting future direction.

## Materials and methods

### Plant materials and growth conditions

The single *ros1-7* mutant used in this study is in the Columbia-0 (Col-0) background (Williams et al. 2015). The *dme*, and *drdd* (*dme-2; ros1-3; dml2; dml3*) used in this study are derived from a cross between *dme-2* in the Columbia *glabrous* (Col-gl) background and *rdd* (*ros1-3; dml2; dml3,* a majority Col-0 background with T-DNAs backcrossed from Ws-0 (Penterman et al. 2007)) expressing *pAGL61::DME* (Williams et al. 2022). The *rdd* mutants used in this study are progeny of the original mutant line (Penterman et al. 2007) and are not expressing *pAGL61::DME*. Plants were grown in long-day conditions (16 h light / 8 h dark) at 22°C and 60% humidity in a Percival AR100L3, or a Conviron PGR15 growth chamber. Plants used for regeneration assays were grown in sterile conditions on 0.5 x MS medium with 0.8% agar, pH 5.8. After excision, regenerating explants were incubated in a Percival CU-36L5 tissue culture chamber in long-day conditions (16 h light / 8 h dark) at 22°C and 70% humidity with a light level of 90 µmol m□² s□¹. Quadruple homozygous mutants *dme;ros1;dml2;dml3* expressing *pAGL61::DME* (termed *drdd (Williams et al. 2022)*) were genotyped by PCR. The primers used for each mutant and WT allele are listed in Table S1. Wild-type segregants from the cross used to generate *drdd* were used as a closely related WT control for *drdd*.

### Regeneration Assays

For all regeneration assays described below, media are defined as follows:

1. Callus induction medium (CIM): Gamborg B5 medium containing Gelzan (0.25% w/v), glucose (2% w/v), 2,4-D (0.5 mg/liter), and kinetin (0.1 mg/liter), pH 5.8.
2. Shoot induction medium (SIM): Gamborg B5 medium containing Gelzan (0.25% w/v), glucose (2% w/v), indole-3-acetic acid (0.15 mg/liter), and 2-iPA (0.5 mg/liter), pH 5.8.
3. Hormone-free B5: Gamborg B5 medium containing Gelzan (0.25% w/v), glucose (2% w/v).

For all assays listed below, explants were inspected and imaged using a Leica M205 FA microscope with a Leica DFC7000 T camera, the number of leaves on each explant was manually assessed under a dissecting microscope, and the regeneration area of explants was quantified with Fiji (https://imagej.net/Fiji).

#### Hypocotyl CIM > SIM assay

*Arabidopsis* seeds were sterilized and stratified for 3 days before being grown on 0.5 x MS medium in the dark for 3 days to promote etiolation. Well-elongated seedlings were selected, excised using a disposable scalpel and transferred to CIM for one week, followed by SIM for 4 weeks under the same growth conditions. Results from each assay can be found in Table S2.

#### Root tip CIM > SIM assay

*Arabidopsis* seeds were sterilized and stratified for 3 days before being grown on vertical plates containing 0.5 x MS medium for 7 days under long-day conditions. Root tip explants were cut from the primary root tip and transferred to CIM for 4 days followed by SIM for 3 weeks.

#### Hypocotyl CIM > Hormone-free B5 assay

*Arabidopsis* seeds were sterilized and stratified for 3 days before being grown on 0.5 x MS medium in the dark for 7 days to promote etiolation. Etiolated hypocotyls were excised and transferred to CIM for 9 days, followed by hormone-free B5 for 13 days.

#### Hypocotyl > SIM assay

*Arabidopsis* seeds were sterilized and stratified for 3 days before being grown on 0.5 x MS medium in the dark for 7 days to promote etiolation. Etiolated hypocotyls were excised and placed directly on SIM for 21 days.

#### Hypocotyl Hormone-free B5 root regeneration

*Arabidopsis* seeds were sterilized and stratified for 3 days before being grown on 0.5 x MS medium in the dark for 7 days to promote etiolation. Etiolated hypocotyls were excised and placed directly on hormone-free B5 for 19 days.

#### First true leaf hormone-free B5 assay

*Arabidopsis* seeds were sterilized and stratified for three days at 4°C, then grown for 21 days on 0.5 x MS medium in Magenta tissue culture boxes. First true leaves were then excised at the base of the leaf blade (fully omitting the petiole) and placed on hormone-free B5. Explants on hormone-free B5 were incubated for 6-8 weeks and manually inspected for the appearance of regenerated shoots. The frequency of shoot regeneration was calculated from the subset (approximately 50%) of leaf cuttings that successfully regenerated roots (Figure S2).

### Enzymatic Methyl-seq

Three to four first true leaves were collected from plants 21 days after stratification (with the exception of the postR_0_ samples, which were generated from single rosette leaf samples after regeneration), and DNA was extracted using a CTAB protocol. Enzymatic methyl-seq (Feng et al. 2020) was performed using the NEBNext EM-seq kit (#E7120) using 25-50ng of DNA as input with 8 cycles of amplification. Libraries were quantified using a KAPA Library Quant Kit according to the manufacturer’s instructions, and quality was verified using an Agilent Fragment Analyzer. Samples were sequenced using an Illumina NovaSeq X (150 bp, paired-end reads). Library fragment analysis and sequencing were performed by the University of California, Berkeley QB3 Genomics facility. The final sequencing output for each sample is listed in Table S3.

### Whole-genome DNA methylation analysis

Mapping of whole-genome bisulfite sequencing and EM-seq data was performed as described (Williams et al. 2023). Briefly, reads were preprocessed with Trim Galore v0.6.6 (Babraham Bioinformatics) to remove adaptors. Bismark v0.22.3 (Krueger and Andrews 2011) was used to map filtered reads to the *A. thaliana* TAIR10 genome, allowing one mismatch per read in the seed region and removing PCR duplicates. Methylation values for each cytosine were calculated as #C/(#C+#T) using the Bismark methylation extractor function. The efficiency of EM-seq conversion was verified by quantifying the percentage of methylation for reads mapped to the chloroplast.

Differentially methylated regions (DMRs) were called using the R package DMRcaller v1.36.0 (Catoni et al. 2018). Briefly, the genome was divided into 300 bp bins containing at least 10 cytosines with >= 5 reads coverage per cytosine. Filtered bins with an absolute CG methylation difference of at least 0.35 and cutoffs of Benjamini–Hochberg corrected false discovery rate < 0.01 were computed as DMRs. For CHG, the cutoff was .2, and for CHH, the cutoff was .15. Bins were merged into larger windows, provided that the merged window remained statistically significant. DMRs were called between WT and preR or postR *drdd*, as well as between preR *drdd* and postR_0_ or postR_1_ lines. In defining DNA methylation changes linked to regeneration in postR *drdd* lines, DMRs between WT and preR *drdd* were filtered out, as these were assumed to represent previously described methylation changes linked to the loss of DRDD activity and unrelated to propagation by regeneration. The remaining regions, termed R-DMRs, were then analyzed. R-DMRs that were also called as differentially methylated between preR *drdd* and postR were categorized as “Group I de novo”, and the remaining R-DMRs were categorized as “Group II exacerbated”. Additionally, R-DMRs were assigned as shared between multiple lines or unique to individual lines using bedtools intersect (Quinlan and Hall 2010). DMR coordinates can be found in Datasets S2, S3, and S4. In Figure S3, DMRs were summed across 350-kb windows. For Figures 3C and 3E, methylation levels were averaged across 50-bp windows overlapping DMRs and extending 300 bp upstream and downstream of DMR boundaries. The heatmaps in Figures 3C and 3E were generated in R using the package ComplexHeatmap (Gu et al. 2016), using hierarchical clustering to sort DMRs.

The distance from Araport11-annotated transcription start sites (TSSs) to the closest DMR (Fig. 5A) was calculated using bedtools closest. Control windows were generated by selecting an equivalent number of random windows (whereby n = the number of DMRs for each sequence context) from 300-bp windows, with windows of high methylation (>65% CG, >80% CHG, and >85% CHH) filtered out, to reflect regions statistically eligible to be called as a DMR. Significance was calculated between the distance distribution of R-DMRs and random control windows using a two-sided K-S test.

In Figure 5E, the coordinates of differentially methylated regions proximal to regeneration-associated genes were selected from within 2.5 kb of the start and end sites of regeneration-associated genes and non-overlapping with other genes. Exact coordinates were selected by hand after inspecting methylation profiles of each locus and can be found in Table S3. 31 regeneration-associated genes possessed methylation increases in postR *drdd* vs preR. In generating the heatmap shown in Figure 5E, methylation levels were averaged across DMR regions and the heatmap was generated using the ComplexHeatmap package. CMT2-type and RdDM-type CHH DMR regions were selected on the basis of CMT2 preference for the CAA and CTA subcontexts (Gouil and Baulcombe 2016). Briefly, CHH cytosines with greater than 5 reads were selected. Regions were set as CHH DMRs that gained methylation. For each DMR, the methylation level for every possible CHH subcontext was calculated. Regions were then selected as putative CMT2 targets if the difference between the averaged CAA/CTA methylation level and the remaining 7 subcontexts was greater than 8%. Regions were selected as putative RdDM target regions if the total % mCHH was greater than 2%, and the difference between the averaged CAA/CTA methylation level and the remaining 7 subcontexts was less than 5%. The remaining regions were labeled as “intermediate”.

### RNA sequencing and analysis

For RNA sequencing, 3 to 4 first true leaves per biological replicate were collected from plants grown on soil at 21 days post stratification. Seven biological replicates per genotype/treatment were used for all RNA-seq experiments. Total RNA was isolated using a QIAGEN RNeasy plant mini kit according to manufacturer’s instructions. Non-stranded libraries for poly-A-tailed RNA were generated by Novogene and sequenced using 150-bp paired end protocol (also performed by Novogene). Mapping was performed as follows: adapters were trimmed using cutadapt v5.2 (Martin 2011) and Trim Galore v0.6.6 (Babraham Bioinformatics) by trimming 9 bp from the 5’ end of reads and enforcing a 3’ end quality of > 25%. Reads were mapped to the Araport11 genome using STAR v2.7.1 (Dobin et al. 2013), permitting 0.05 mismatches as a fraction of total read length and discarding reads that did not map uniquely. The final sequencing output for each sample is listed in Table S4. Normalization of read counts was performed using HTSeq v2.0 (Anders et al. 2015) and DESeq2 (Love et al. 2014) version 1.40.1 was used to call DEGs based on a minimum of two-fold change in expression and a Benjamini–Hochberg corrected P value < 0.05. DESeq output can be found in Datasets S1, S5, S6, and S7. Heatmaps, correlation, and PCA analyses were generated using R.

### Floral bud histology

Tissues were fixed in 10 mM Sørensen’s buffer pH 7.2 using a microwave as described (Schichnes et al. 2001). Samples were dehydrated using a graded ethanol series, transferred to isopropanol, then embedded in paraffin. Sectioning was performed using a Leica RM2255 microtome. Slides were deparaffinized in xylene, then rehydrated through a graded ethanol series and stained with Safranin-O, differentiated with picric acid, and stained with Fast Green, according to Johansen’s Safranin and Fast Green staining protocol (Ruzin 1999). Slides were imaged with a Zeiss AxioImager M1.

## Acknowledgements

We are grateful to Denise Schichnes for assistance with floral bud histology and imaging. AI was used to assist with manuscript proofreading only.

## Author Information

## Contributions

NKS, YZ, RMH and BPW performed experiments. NKS, YZ and BPW designed experiments. NKS and BPW conceived of the study and wrote the paper.

## Ethics Declarations

Competing Interests:BPW and NKS are inventors on a pending patent application filed by The Regents of the University of California relating to methods for plant regeneration without callus induction (International Patent Application No. PCT/US2024/052027; publication WO2025085785A2). The other authors declare no competing interests.

---

**Supplementary Figure 1:**
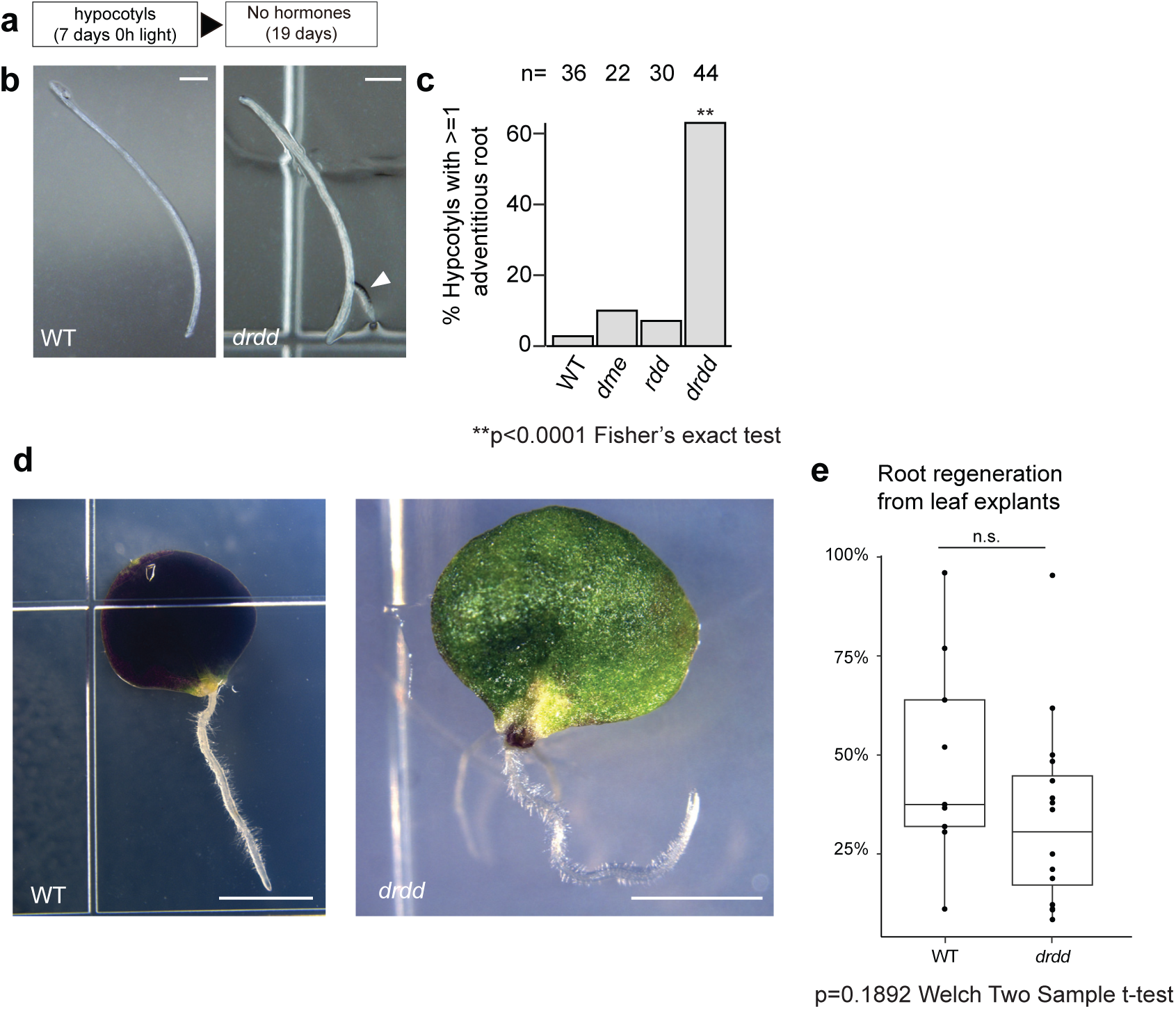
Hormone-free de novo root regeneration in hypocotyl and leaf explants. A) Description of experimental regimen. B) Representative images of hypocotyls after 19 days post excision on hormone-free media. White arrow: DNRR. C) Quantification of B). D) Representative images of leaf explants incubated on hormone-free media showing adventitious root regeneration. E) Quantification of D).

**Supplementary Figure 2:**
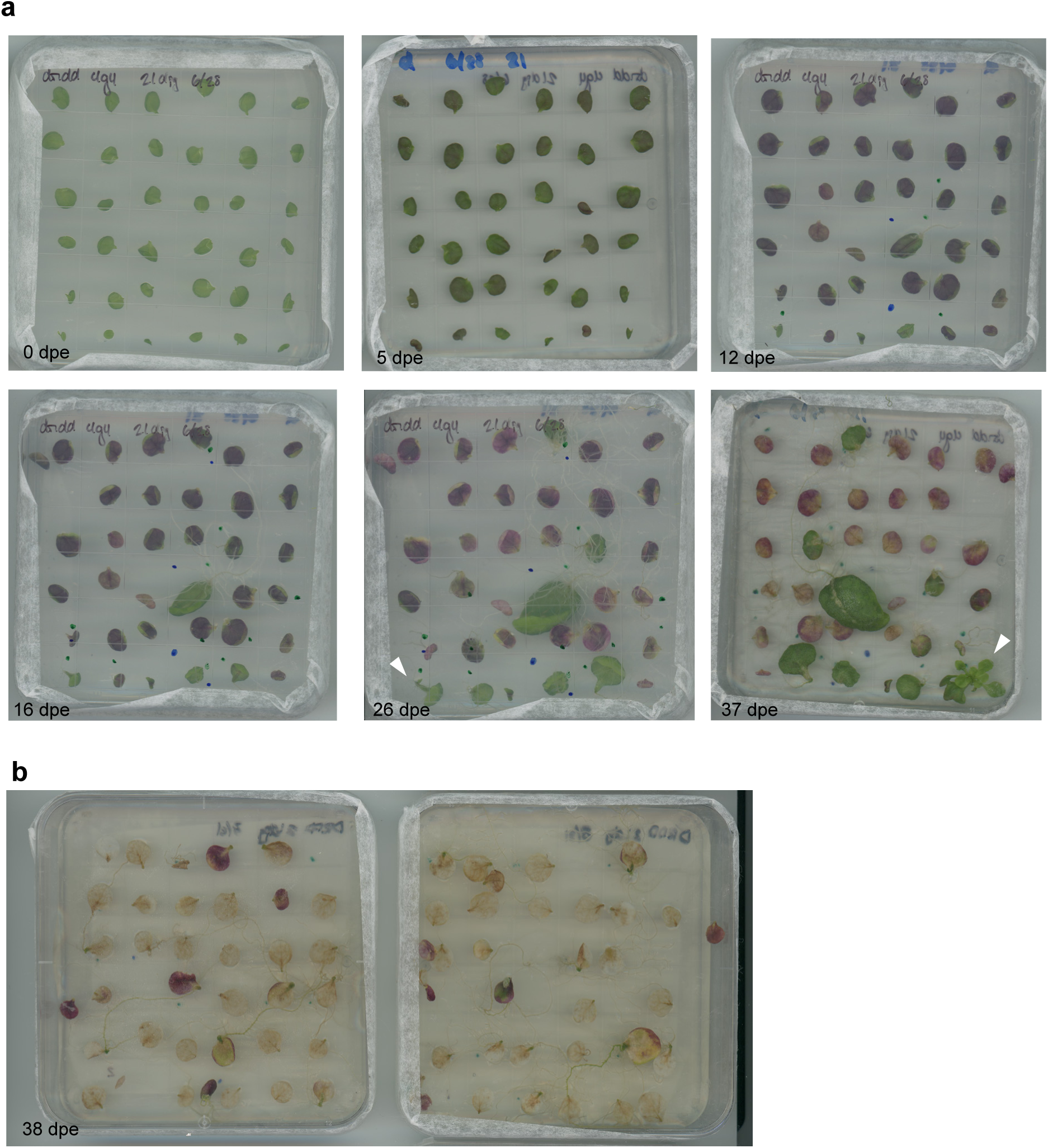
Whole plate images of leaf explant assay. A) Time course scans of one plate of leaf explants. Explants are from preR *drdd* plants. Note that some scans show the abaxial side and some show the adaxial side of leaves. White arrows: DNSR. B) Representative images of DRDD explants at 38 days post excision.

**Supplementary Figure 3:**
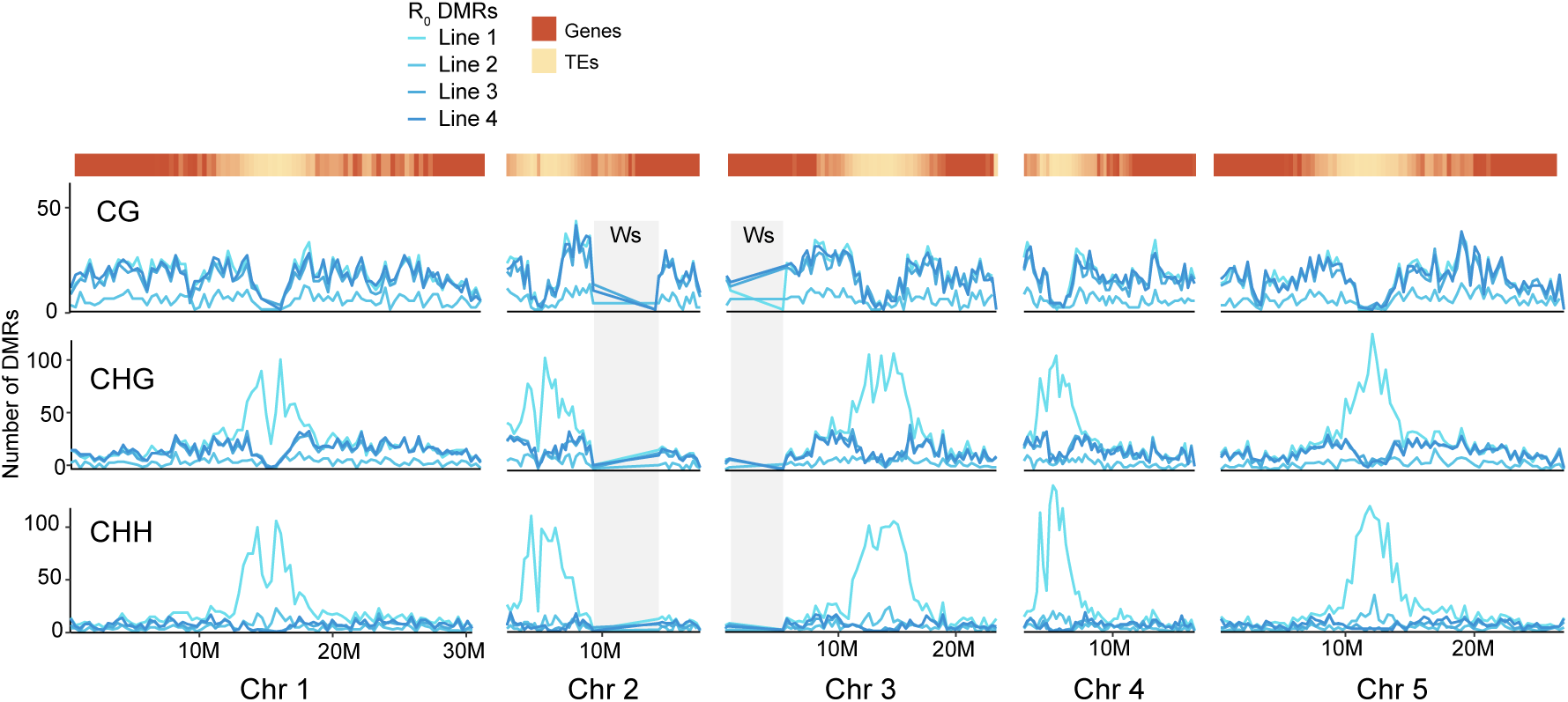
Distribution of DMRs along chromosomes in all regenerated lines. Regions inherited from Ws that were excluded from analysis are indicated by grey boxes. Enrichment of TEs relative to genes is used to indicate centromeric regions.

**Supplementary Figure 4:**
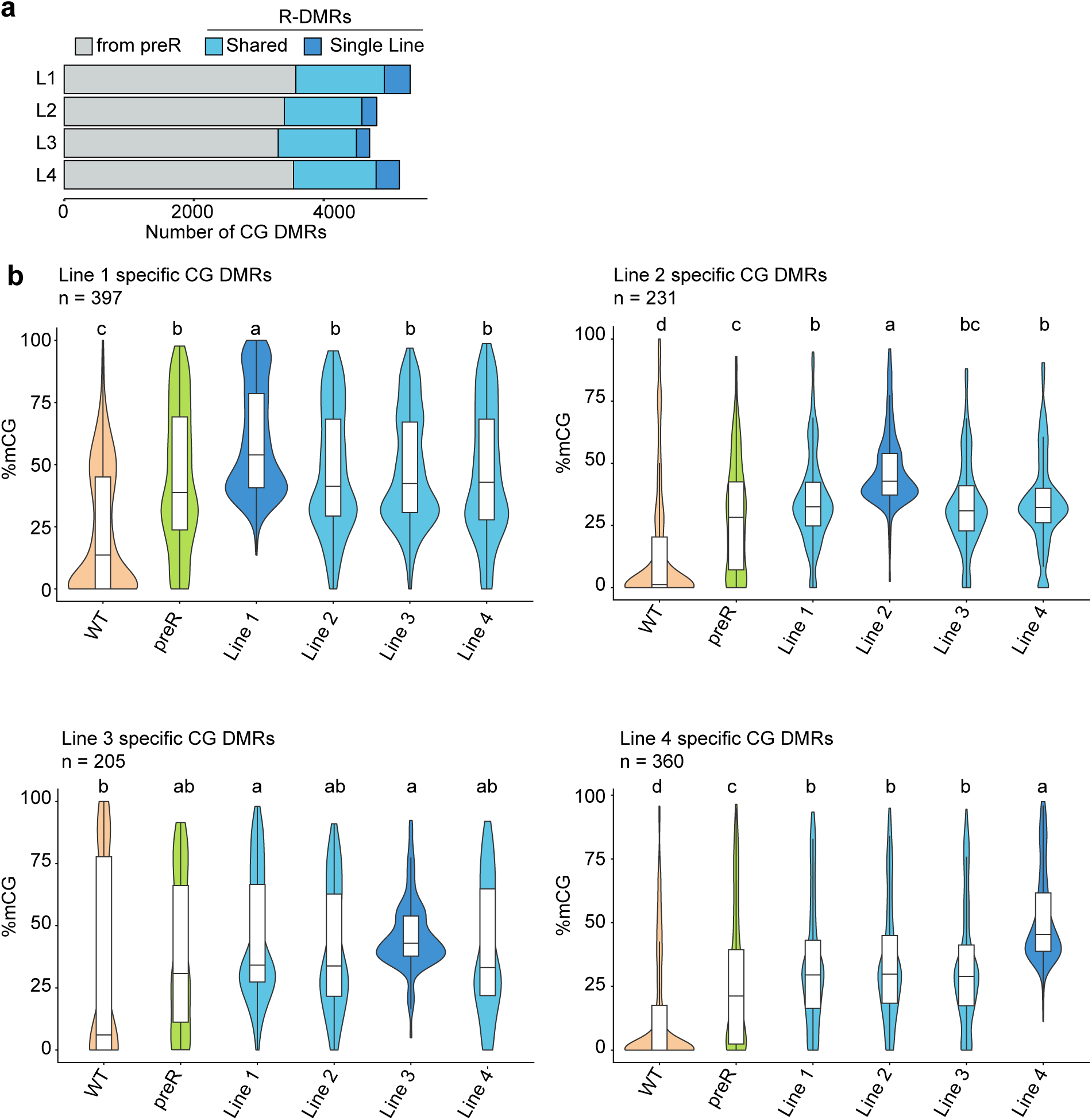
Quantification of methylation at single line DMRs. A) All DMRs in postR plants were separated into categories of “called in preR”, “shared” (the same region was called in at least 2 postR lines), or “single line” (the DMR was called in only that line). B) CG methylation level in each line at DMRs called in only a single line; line in which regions were called is indicated in darker blue. Box plot indicates median, 25^th^ and 75^th^ percentile; whiskers represent 1.5x the inter quartile range. Outliers not shown. One way ANOVA calculated with p < 0.05.

**Supplementary Figure 5:**
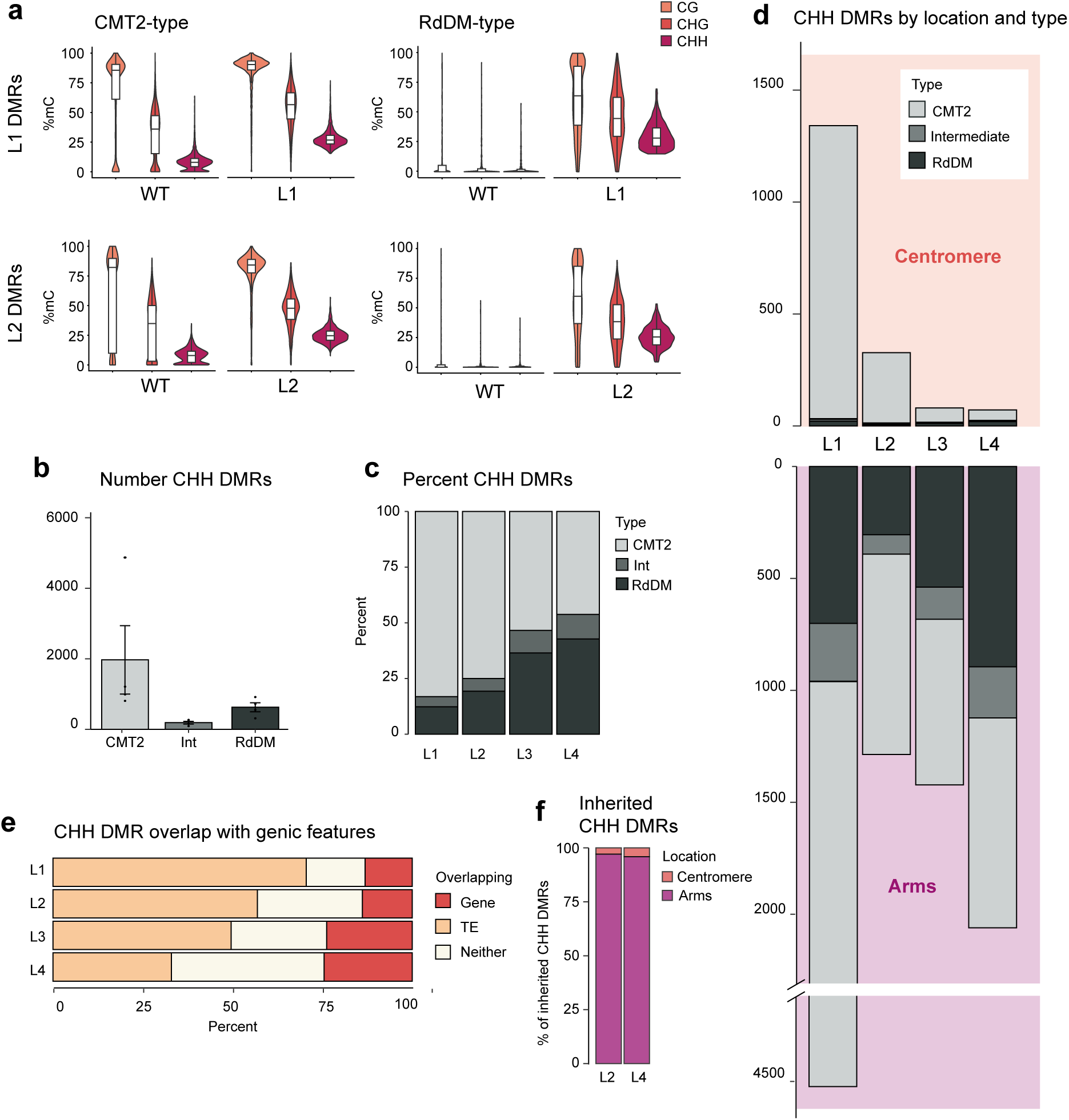
CHH DMRs can be sorted into CMT2-maintained and de novo RdDM-established. A) Methylation level in WT and respective postR_0_ line at regions called as CMT2-type and RdDM-type in Line 1 and Line 2 postR_0_ CHH DMRs. B) Number of each type of CHH DMR in each postR_0_ line. C) Percent of CHH DMRs that fall into each category in each postR_0_ line. D) Location of CHH DMRs by type in each line. E) Overlap of CHH DMRs with genes and TEs. F) Percent of each type of CHH DMR that was called in the respective postR_1_ line.

**Supplementary Figure 6:**
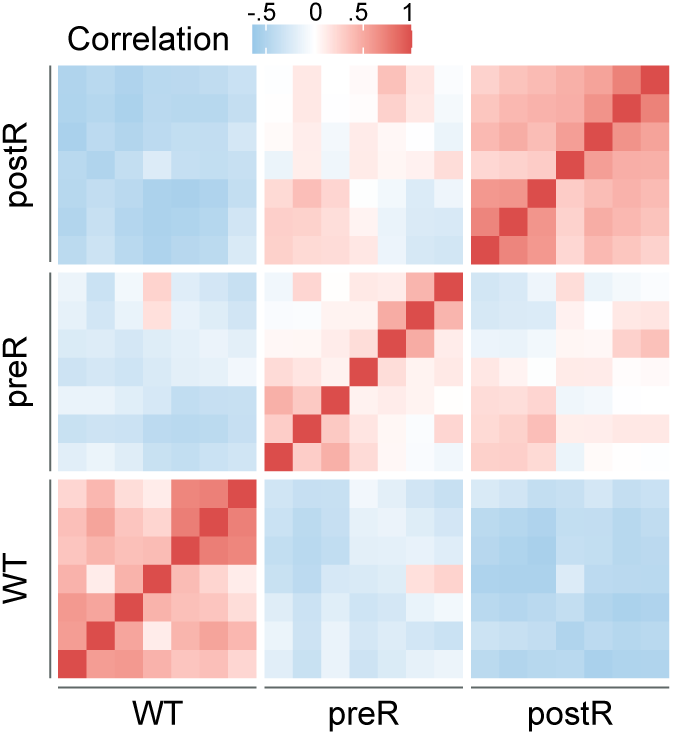
Heatmap denoting Pearson’s correlation coefficient of expression levels of all differentially expressed genes (combined from preR vs WT and postR_1_ vs WT) between each pairwise comparison of individual RNA-seq samples.

## Notes

### Competing Interest Statement

BPW and NKS are inventors on a pending patent application relating to methods for plant regeneration without callus induction (International Patent Application No. PCT/US2024/052027; publication WO2025085785A2). The other authors declare no competing interests.

### Summary of Updates

Genomics analyses and data presentation revised for more clarity. Main conclusions are unchanged

### Funding

- Bakar Bioenginuity Hub

